# A neuroimaging dataset during sequential color qualia similarity judgments with and without reports

**DOI:** 10.1101/2024.05.16.594267

**Authors:** Takahiro Hirao, Mitsuhiro Miyamae, Daisuke Matsuyoshi, Ryuto Inoue, Yuhei Takado, Takayuki Obata, Makoto Higuchi, Naotsugu Tsuchiya, Makiko Yamada

## Abstract

Recent neuroscientific research has advanced our understanding of consciousness, yet the connection between specific qualitative aspects of consciousness, known as “qualia,” and particular brain regions or networks remains elusive. Traditional methods that rely on verbal descriptions from participants pose challenges in neuroimaging studies. To address this, our group has introduced a novel “qualia structure” paradigm that leverages exhaustive, structural, and relational comparisons among qualia instead of verbal reports. In this study, we present the first fMRI dataset that captures relational similarity judgments among two out of nine color qualia per trial from 35 participants. This dataset also includes a “no-report” condition in half of the trials to assess the impact of overt reporting. Additionally, each participant’s color discriminability was evaluated with a hue test conducted outside the scanner. Our data offer valuable insights into the brain functions associated with color qualia and contribute to a deeper understanding of the neural foundations of consciousness.

## Background & Summary

One of the biggest mysteries in current science is the relationship between our subjective experiences and the brain. How are our qualitative aspects of experience, such as redness of red, supported by physical substrates in the brain? Some philosophers have suggested that scientific methods cannot be applied to study the qualitative aspects of consciousness, or qualia, for they are deemed intrinsic, ineffable and private (D. C. Dennett, 1988). While direct description of a quale remains elusive, other philosophers and scientists have found a path forward in characterizing them through exhaustive structural and relational comparisons among qualia (D, Rosenthal, 2015; S. B. Fink, et al., 2021; N. Tsuchiya and H. Saigo, 2021; J. Kleiner, 2020; D. J. Chalmers, 1996; T. Nagel, 1974; N. Tsuchiya et al., 2022).

Within this pursuit, the examination of qualia through the lens of similarity judgment, particularly in the domain of color, has proven to be a potent tool within the realm of behavioral psychophysics (C. E. Helm, 1964; R. N. Shepard and L. A. Cooper, 1992; G. P. Epping et al., 2023; A. M. Zeleznikow-Johnston, 2023; T. Indow, 1988; G. V. Paramei et al., 2001; A. Boehm et al., 2014; G. Kawakita et al., 2023; J. M. Boesten,2005). Yet, the exploration of the neural underpinnings of such “similarity judgements” among color qualia has remained a largely uncharted territory.

Evidence from clinical studies has linked the neural basis of the color qualia to specific brain regions, namely, visual areas V4 and V8, within the ventral occipitotemporal cortex. The impairment of these areas has been correlated with subtle yet discernible declines in color processing abilities in affected patients (L. M. Vaina, 1994; S. E. Bouvier and S. A. Engel, 2006).

Further insights have emerged from functional magnetic resonance imaging (fMRI) investigations in healthy individuals, implicating V4/V8 in color qualia processing (N. Hadjikhani et al., 1998; S. Zeki et al., 1991; A. A. Brewer et al., 2005). While the precise roles of these regions continue to be a subject of ongoing debate (A. W. Roe et al., 2012), recent research trends converge on the idea that color qualia is intricately supported by neural mechanisms along the ventral surface of the occipitotemporal cortex (M. S. Beauchamp et al., 1999; R. Shapley and M. Hawken, 2011).

Expanding this body of fMRI research, Brouwer and Heeger (2009) and (2013) introduced a more nuanced approach, delving into the relational characterization of neural activity related to color qualia structures. They departed from the conventional focus on overall V4/V8 activity and instead employed multi-voxel pattern analyses, encompassing decoding, reconstructing, and clustering. This methodology extended to the magnetoencephalography (MEG) to pinpoint the precise temporal dynamics of these neural interactions (I. Rosenthal et al., 2021). This line of research holds significant promise in bridging the gap between the structural characteristics of color qualia unveiled through psychophysics (C. E. Helm, 1964; R. N. Shepard and L. A. Cooper, 1992; G. P. Epping et al., 2023; A. M. Zeleznikow-Johnston, 2023; T. Indow, 1988; G. V. Paramei et al., 2001; A. Boehm et al., 2014; G. Kawakita et al., 2023; J. M. Boesten,2005) and the structural characteristics of brain activity patterns and connectivity.

Here, we provide a novel fMRI dataset poised to enrich existing literature in three keyways. First, unlike previous studies (G. J. Brouwer et al., 2009, 2013; I. A. Rosenthal 2021], our fMRI data was acquired while participants engaged in similarity judgments between two color patches (See Methods). This departure from psychophysics measurements outside the scanner is significant, as acquiring similarity judgments inside the scanner is highly time consuming and expensive. Second, to disentangle the neural substrates of color qualia from those associated with reporting similarity judgements, we implemented a no-report paradigm (N. Tsuchiya, 2015) (See Methods). This approach is crucial given the growing body of neuroimaging evidence implicating regions like the prefrontal cortex in the act of reporting (fMRI: A. A. Farooqui, 2016; S. Frässle, 2014; T. Dellert, 2021, M/EEG: M. A. Cohen et al., 2020; C. Sergent et al., 2021; J. P. Shafto and M. A. Pitts, 2015; E. Rowe, 2023). By combining data from both reporting and no-reporting trials, our dataset offers a unique opportunity to isolate the neural correlates of color qualia structure from those linked to the reporting of the similarity judgments. Third, we present a comprehensive behavioral characterization (outside scanner) of hue tests for each participant, enabling further exploration of the neural basis of the individual differences in color qualia and similarity judgments.

## Methods

### Participants

Thirty-five adults (17 male and 18 female participants, aged 21–59 years [*M* ± *SD*: 40.6 ± 12.5 years]) were paid to participate in the study. We recruited the participants using the volunteer recruiting system of the National Institutes for Quantum Science and Technology (Table 1). All participants had no history of neurological or psychiatric disorders and did not take any medications that might interfere with task performance or fMRI data. We confirmed that they did not have red-green color deficiencies using the 38-plates-version of the Ishihara test (S. Ishihara, 1990). Due to excessive head motion, we excluded four participants (see Technical Validation). The study was approved by the Ethics Committee of the National Institutes for Quantum Science and Technology in Chiba, Japan, and all participants provided written informed consent.

### Experimental procedure

After completing the color vision assessments (ND-100 Hue test, Ishihara test) and the questionnaires (the Edinburgh Handedness Inventory, the State-Trait Anxiety Inventory, Beck Depression Inventory-II), participants practiced the color similarity task outside the scanner first, then inside the scanner. Then, they underwent the MRI scanning (Figure 1a).

**Figure 1.**
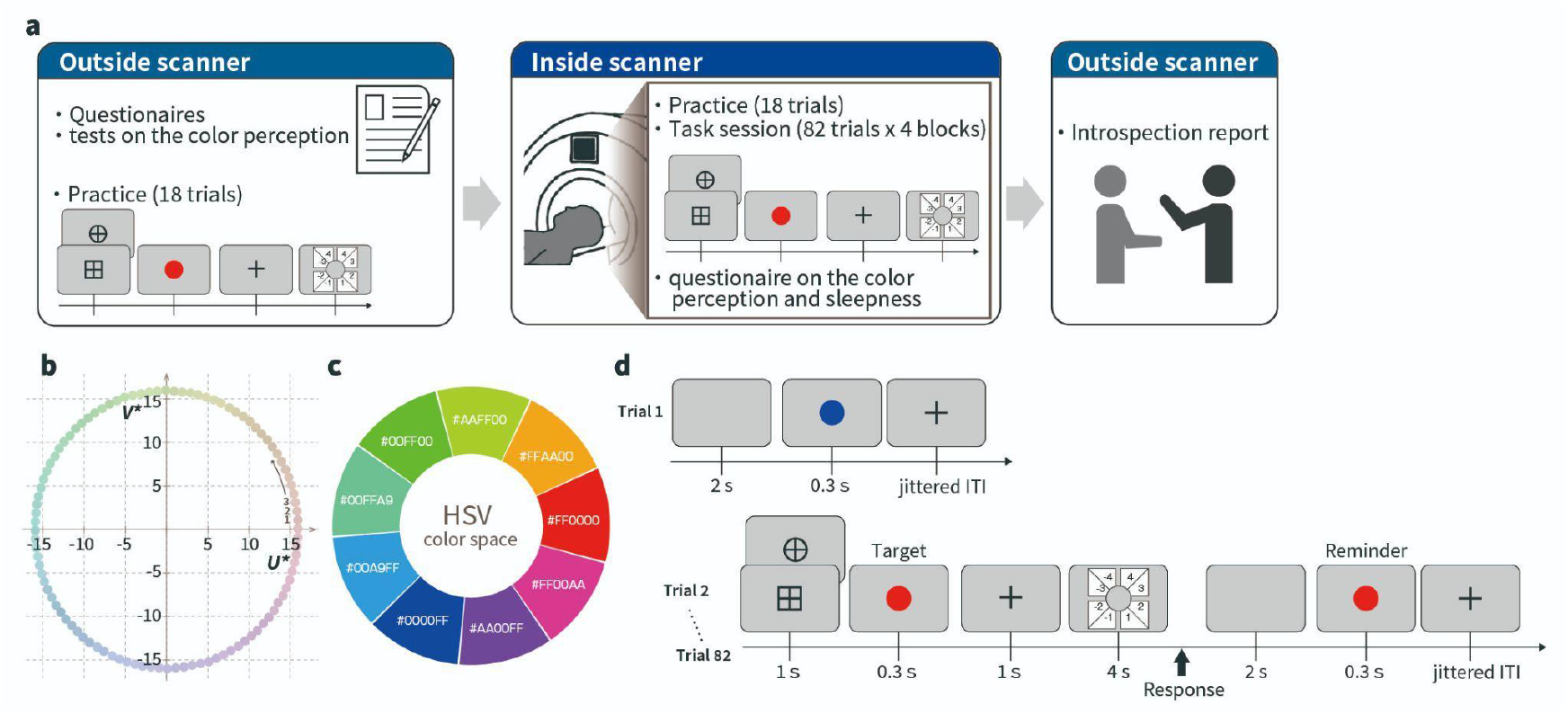
Overview of the experiment. (a) Outline of the experimental workflow used to acquire the dataset. (b) Image for the 100 colors of the ND-100 hue test. The 100 colors on the chromaticity based on the CIE 1964 color space were depicted on the grid. Each color is separated by one color difference [a color difference is defined by a national bureau of standards (NBS) unit]. (c) The nine colors used in the fMRI task. (d) Illustration of the fMRI task procedure.

### Color vision assessments

We assessed participants’ color vision with the Ishihara test and the ND-100 Hue test. We used the Ishihara test to assess color vision deficiency with 38 plates (the full version).

We also used the ND-100 Hue test to evaluate their color recognition capability. The ND-100 hue test, developed by the Japan Color Research Institute (Japan Color Enterprise Co., Ltd., 1972), uses 100 hues that are equally spaced on a circumference that is centered on the standard illuminant C and has a length of 100 color difference units on the CIE 1964 uniform color space with a brightness of 6 (Figure 1b). Participants were asked to sort 25 hues randomly arranged in one row into the correct gradient order within a time limit of 2 min. A total of four rows of sorting was completed. The 100 hues were assigned a natural number from 1 in an ascending order along the increase of the polar angle (Figure 1b). Thus, the difference in the assigned number between the two neighboring hues is 1. For each hue, we calculated the absolute value of the difference between the numbers of each hue’s two neighboring hues, and subtracted 2 from the sum of the two values. We take this as the discrimination score for that hue, which is zero if the hue is correctly arranged by the participant. Table 1 (Column: ND-100 test) lists the sum of the discrimination scores across 100 hues for each participant as a summary measure of color recognition capability.

**Table 1.** Demographic information and results of questionnaires for all participants in this study. *STAI* State-Trait Anxiety Inventory, *BDI-II* Beck Depression Inventory-II.

| ID | Sex | Age | Color vision assessments |  | Questionnaires |  |  |  | fMRI Task |
| --- | --- | --- | --- | --- | --- | --- | --- | --- | --- |
|  |  |  | Ishihara test | ND-100 hue test | Edinburgh Handedness Inventory | State | Trait | BDI-II | Cue for report trial |
| 1 | F | 32 | 2 | 228 | 90.0 | 22 | 24 | 10 | Square |
| 2 | M | 33 | 1 | 60 | 100.0 | 44 | 53 | 11 | Square |
| 3 | M | 50 | 1 | 96 | 87.5 | 32 | 36 | 3 | Circle |
| 4 | F | 43 | 1 | 40 | 100.0 | 45 | 52 | 13 | Square |
| 5 | M | 59 | 0 | 164 | 90.0 | 35 | 38 | 9 | Square |
| 6 | F | 56 | 4 | 228 | 100.0 | 40 | 39 | 8 | Circle |
| 7 | M | 37 | 2 | 68 | 78.9 | 33 | 52 | 5 | Square |
| 8 | F | 47 | 1 | 68 | 100.0 | 30 | 31 | 6 | Square |
| 9 | F | 50 | 1 | 120 | 100.0 | 44 | 49 | 20 | Circle |
| 10 | F | 21 | 1 | 88 | -20.0 | 23 | 39 | 5 | Circle |
| 11 | M | 24 | 2 | 148 | 100.0 | 39 | 42 | 12 | Circle |
| 12 | F | 38 | 0 | 104 | 47.4 | 21 | 25 | 0 | Square |
| 13 | M | 25 | 0 | 40 | 5.9 | 49 | 60 | 5 | Circle |
| 14 | M | 22 | 3 | 500 | 100.0 | 47 | 65 | 24 | Square |
| 15 | F | 57 | 1 | 76 | 90.0 | 35 | 27 | 5 | Circle |
| 16 | F | 21 | 1 | 124 | 100.0 | 47 | 59 | 6 | Circle |
| 17 | F | 36 | 1 | 168 | 100.0 | 37 | 49 | 14 | Square |
| 18 | F | 41 | 0 | 24 | 100.0 | 40 | 51 | 4 | Square |
| 19 | M | 34 | 0 | 152 | 100.0 | 41 | 45 | 2 | Circle |
| 20 | F | 51 | 4 | 160 | 100.0 | 38 | 39 | 7 | Square |
| 21 | F | 55 | 2 | 104 | 100.0 | 35 | 55 | 20 | Circle |
| 22 | F | 42 | 4 | 104 | 100.0 | 44 | 40 | 2 | Circle |
| 23 | M | 25 | 0 | 68 | 100.0 | 44 | 43 | 4 | Circle |
| 24 | M | 55 | 0 | 104 | 100.0 | 35 | 43 | 4 | Square |
| 25 | M | 46 | 1 | 64 | 100.0 | 41 | 33 | 0 | Circle |
| 26 | F | 54 | 3 | 180 | 100.0 | 36 | 49 | 22 | Square |
| 27 | F | 21 | 0 | 96 | 78.9 | 36 | 46 | 6 | Circle |
| 28 | M | 53 | 0 | 152 | 76.5 | 40 | 58 | 23 | Circle |
| 29 | F | 43 | 1 | 48 | 100.0 | 30 | 42 | 3 | Circle |
| 30 | M | 54 | 1 | 144 | 89.5 | 36 | 37 | 1 | Square |
| 31 | M | 43 | 0 | 48 | 100.0 | 43 | 43 | 7 | Square |
| 32 | M | 57 | 1 | 116 | 70.0 | 44 | 42 | 9 | Square |
| 33 | M | 42 | 4 | 168 | 100.0 | 24 | 23 | 0 | Circle |
| 34 | F | 30 | 0 | 16 | 78.9 | 32 | 48 | 6 | Square |
| 35 | M | 24 | 1 | 100 | 100.0 | 28 | 31 | 1 | Square |

### Questionnaires

The Edinburgh Handedness Inventory was used to assess handedness (R. C. Oldfield, 1971). The laterality quotient was calculated; positive values indicate a preference for using the right hand, and negative values indicate a preference for using the left hand. The State-Trait Anxiety Inventory (C. D. Spielberger et al., 1971; 1983) and Beck Depression Inventory-II(A. Beck, 1996; M. Kojima et al., 2002) were used to measure the levels of anxiety and depression, respectively. High scores on both scales indicate severe symptomatic states.

One-back color similarity judgment task with report/no-report instructions in the fMRI: Participants performed a color similarity rating task with nine colors. The color calibration of the monitors was performed to output the intended colors. In the color calibration, a color and luminance meter was used to determine the sRGB values necessary to represent the nine colors described above (Strokes et al., 1996). The hue of nine colors were 0, 40, 80, 120, 160, 200, 240, 280, and 320 degrees in the HSV color space (counterclockwise from #FF0000) (Figure 1c). All nine colors had a 100% saturation and a 100 % value. We presented the hue stimuli on a gray background (0, 0, and 50% in the HSV color space). The task was programmed using presentation software (Neurobehavioral Systems, Inc., Berkeley, CA, USA).

Each trial starts with a 1,000-ms cue (overlaid on a fixation) instructing participants to report (circle) or not to report (square) color similarity. The comparison between the report and no-report conditions allowed us to estimate the neural activity associated with the act of reports (and preparation of reports, including explicit judgments and memorization of color qualia) (eg, N. Tsuchiya, 2015). Following the cue, we presented a 300-ms target color, a 1,000-ms fixation, a 4,000-ms response display, a 2,000-ms blank, and a 300-ms color patch. The last color patch was the same color as the current target and presented again as a reminder for comparison with the next trial. We asked participants to rate the similarity level on an eight-point scale of -4, -3, -2, -1, 1, 2, 3, and 4. Negative and positive numbers represent “dissimilar” and “similar” responses. Higher magnitudes represent a more vivid experience. For example, “-4” indicates that two colors were highly dissimilar. The participants made the judgment response by moving the cursor and clicking the button using an MRI-compatible trackball mouse (Current Designs, Philadelphia, USA).

### MRI acquisition

All MRI data were acquired using a Siemens Verio 3 Tesla MRI system with 32 channel head coil (Siemens Healthcare, Erlangen, Germany). Structural images were acquired using a T1-weighted magnetization-prepared rapid gradient-echo (MPRAGE) protocol (repetition time [TR] = 2,300 ms; echo time [TE] = 2.98 ms; flip angle [FA] = 9 °; 176 slices; in-plane field of view = 256 × 240 mm; 1.0 mm isotropic voxel). Functional images were acquired using a multiband, T2*-weighted gradient-echo echo planar imaging sequence (TR = 1,000 ms; TE = 31.6 ms; FA = 61 deg; 45 slices; MB factor = 3; 3.2 mm isotropic voxel; 862 volumes; 14 min 31 sec per 1 run; 4 runs) provided by the University of Minnesota (S. Moeller et al., 2010).

### Technical validation procedure Preprocessing for MRI data

The structural and functional images were preprocessed using Statistical Parametric Mapping software (SPM12: https://www.fil.ion.ucl.ac.uk/spm/software/spm12/) on Matlab R2020b (Mathworks Inc., Massachusetts, USA). We discarded the initial 30 s of functional volumes to ensure a magnetic equilibrium state. First, the functional images were realigned to the first of all images. As in the realignment process, the slice timing was corrected to the first slice. The value for the coregistration of EPIs to a T1 image was estimated, and the segmentation for a T1 image was conducted after coregistration. The EPIs and T1 images were normalized to the standard Montreal Neurological Institute (MNI) space and resliced to a voxel size of 2 mm^3^. Finally, spatial smoothing (Gaussian kernel full-width-half-maximum, FWHM, 4 mm) was applied to the normalized EPIs.

### Estimation of framewise displacement (FD) and temporal signal noise ratio (tSNR)

In order to assess the quality of the data, we performed calculations for FD and tSNR for each functional run utilizing head-motion-corrected data. The assessment of participant head movement involved the computation of FD, relying on the six motion parameters obtained during the realignment process of the fMRI analysis (J. D. Power et al., 2012; 2014). The tSNR for a given functional run was determined by dividing the mean by the standard deviation of the gray matter signals across all voxels in all frames. The calculation of tSNR was conducted on a per-run basis. To facilitate the tSNR computation, the raw DICOM files underwent conversion to 4D NIfTI data using the dcm2niix (https://www.nitrc.org/projects/mricrogl/), followed by realignment and segmentation processing before the tSNR calculation.

### Statistical Analyses

To evaluate time-locked activations, we modeled forty regressors in each run: stimulus presentations for nine target colors, evaluation and nine reminder colors, and responses for the evaluation in both the report and no-report trials. The duration of each regressor was set as 0. Six head motion parameters were treated as nuisance covariates and regressed in the model.

The similarity rating scores obtained from -4 to 4 were converted to dissimilarity scores from 0 (the least dissimilar) to 7 (the most dissimilar) and used for statistical analysis. To check the consistency of rating responses across runs, we averaged dissimilarity scores in each combination of color comparisons on the 1st (run 1-2) and 2nd half (run 3-4) of all runs.

Wilcoxon signed-rank test was used in these comparisons.

## Data Records

All data in this dataset are freely available at the Open Neuro repository (Open Science Framework: https://osf.io/sqd7n/). All files that were used for data acquisition and were produced by the experiment in the fMRI study were put in the “ColorSimilarity_fMRI” folder after personally identifiable information was removed. The “ColorSimilarity_fMRI” folder comprises files for each of the 35 participants (the architecture of these folders is depicted in Figure 2).” There are separate folders for each session within those files, and All DICOM files and behavior files were put in this folder after these files were converted to Brain Imaging Data Structure (BIDS) format version 1.9.0. In the Behavior files, the results of the color comparison in the color similarity rating task conducted in the MRI scanner and the questionnaire results are stored.

**Figure 2.**
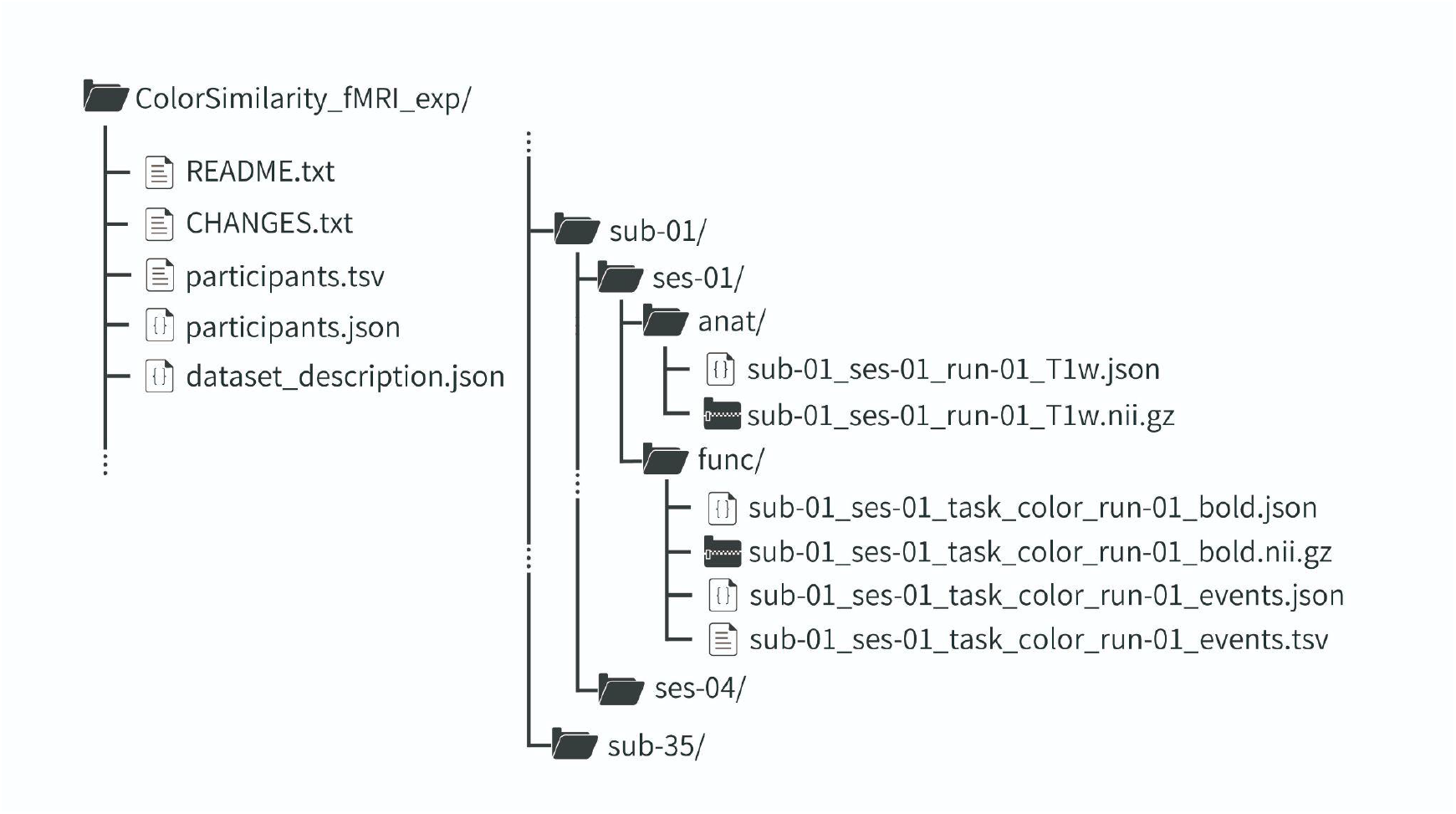
Folder structure for the dataset. The dataset follows the Brain Imaging Data Structure (BIDS) format. The top-level directory “ColorSimilarity_fMRI_exp” contains text files with information about the experiment and participants. Sub-directories are organized hierarchically for 35 participants (e.g., “seb-01”). Within each participant’s directory, there are four session directories which contain anatomical data (“anat”) and functional data (“func”). The functional data includes task-related fMRI data along with event timing information.

## Technical Validation

### 1. Color vision assessment and questionnaires

The results of the color vision assessment and questionnaire were summarized for each participant in Table 1. According to the Ishihara test, no subjects exhibited color blindness values (5 or higher), with an average of 1.3 ± 1.3. The average of the ND-100 Hue test was 119.1 ± 85.1. The mean STAI trait score was 43.1 ± 10.5 and the mean BDI was 7.9 ± 6.8.

### 2. The fMRI task: color similarity rating

#### 2-1. Behavior results

##### 2-1-1. Valid/invalid trials (Table 2)

The number of valid/invalid trials were described in Table2. Moving the cursor and clicking the decision button within 4,000 ms were considered valid for the report trials. Not clicking the decision button within 4,000 ms were considered valid for the no-report trials.

**Table 2.**
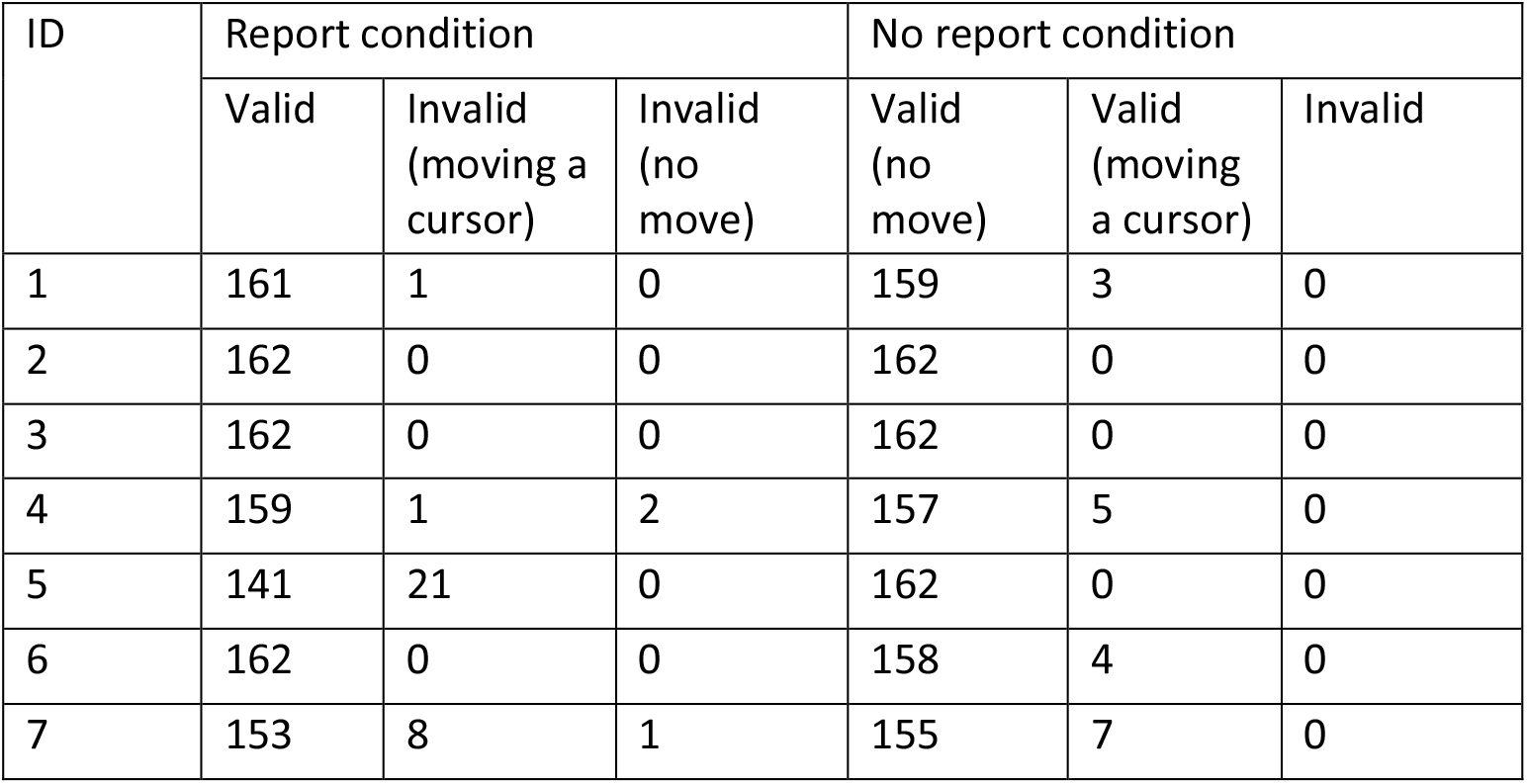
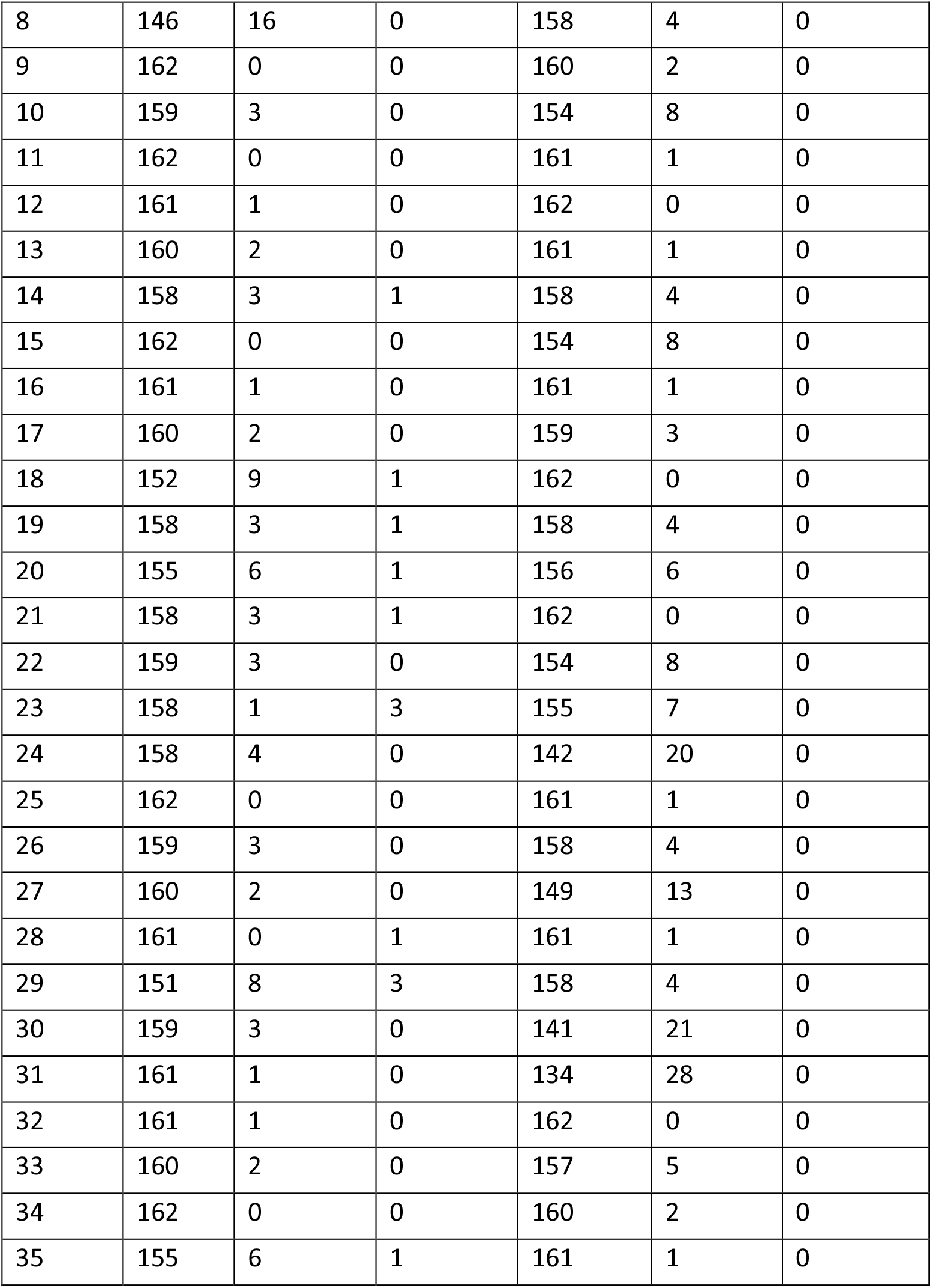
Number of response trials per condition, with the first trial of each run omitted.

##### 2-1-2. Consistency of similarity rating for a particular color pair across runs and participants (Table 3)

To confirm that the similarity judgments in the report trial for a particular color pair were consistent across runs and participants, we compared the similarity values in the first and second halves of the task for each color pair. Wilcoxon signed-rank tests with multiple comparison corrections showed no differences between the first and second halves for each of the 45 color combinations. This result suggests that the responses of the participants were consistent across four runs.

**Table 3.** Grand average of dissimilarity scores for 45 color pairs. This table presents the mean dissimilarity scores for 45 color pairs, derived from run 1-2 and run 3-4. Additionally, it includes corrected p-values obtained through Wilcoxon signed-rank tests with multiple comparison corrections, along with the minimum number of participants available in each case.

| Color1 | Color2 | Dissimilarity score (mean ± sd) | | | Number of available data<br>(Minimum number of participants available) | |
| --- | --- | --- | --- | --- | --- | --- |
|  |  | run1-2 | run3-4 | Corrected p-value | run1-2 | run3-4 |
| #ff0000 | #ff0000 | 0.06 ± 0.23 | 0.00 ± 0.00 | 1.00 | 35 | 33 |
| #ff0000 | #ffaa00 | 2.24 ± 1.02 | 2.24 ± 0.97 | 1.00 | 35 | 35 |
| #ff0000 | #aaff00 | 5.31 ± 1.2 | 5.74 ± 1.32 | 0.225 | 35 | 35 |
| #ff0000 | #00ff00 | 6.00 ± 1.07 | 6.10 ± 0.95 | 1.00 | 35 | 35 |
| #ff0000 | #00ffa9 | 6.10 ± 0.79 | 6.04 ± 0.77 | 1.00 | 35 | 35 |
| #ff0000 | #00a9ff | 6.27 ± 0.70 | 6.31 ± 0.73 | 1.00 | 35 | 35 |
| #ff0000 | #0000ff | 6.80 ± 0.42 | 6.80 ± 0.47 | 1.00 | 35 | 35 |
| #ff0000 | #aa00ff | 2.69 ± 1.3 | 2.90 ± 1.42 | 0.782 | 35 | 34 |
| #ff0000 | #ff00aa | 1.21 ± 0.65 | 1.39 ± 0.72 | 0.867 | 35 | 35 |
| #ffaa00 | #ffaa00 | 0.06 ± 0.23 | 0.00 ± 0.00 | 1.00 | 35 | 34 |
| #ffaa00 | #aaff00 | 2.51 ± 1.37 | 2.70 ± 1.4 | 0.780 | 35 | 35 |
| #ffaa00 | #00ff00 | 3.99 ± 1.14 | 3.87 ± 1.16 | 1.00 | 35 | 35 |
| #ffaa00 | #00ffa9 | 4.53 ± 1.24 | 4.51 ± 1.17 | 1.00 | 35 | 35 |
| #ffaa00 | #00a9ff | 5.51 ± 0.97 | 5.47 ± 0.93 | 0.804 | 35 | 35 |
| #ffaa00 | #0000ff | 6.39 ± 0.56 | 6.01 ± 0.73 | 0.335 | 35 | 35 |
| #ffaa00 | #aa00ff | 4.81 ± 1.44 | 4.93 ± 1.37 | 1.00 | 35 | 34 |
| #ffaa00 | #ff00aa | 3.31 ± 1.44 | 3.36 ± 1.3 | 1.00 | 35 | 35 |
| #aaff00 | #aaff00 | 0.00 ± 0.00 | 0.17 ± 0.61 | 0.804 | 34 | 35 |
| #aaff00 | #00ff00 | 1.11 ± 0.68 | 1.04 ± 0.68 | 1.00 | 35 | 35 |
| #aaff00 | #00ffa9 | 1.63 ± 0.64 | 1.60 ± 0.85 | 1.00 | 35 | 35 |
| #aaff00 | #00a9ff | 4.26 ± 1.38 | 3.91 ± 1.31 | 0.298 | 35 | 35 |
| #aaff00 | #0000ff | 4.90 ± 1.22 | 5.00 ± 1.36 | 1.00 | 35 | 35 |
| #aaff00 | #aa00ff | 5.36 ± 1.05 | 5.74 ± 0.98 | 0.293 | 35 | 35 |
| #aaff00 | #ff00aa | 5.06 ± 1.24 | 5.47 ± 1.32 | 0.326 | 35 | 35 |
| #00ff00 | #00ff00 | 0.03 ± 0.17 | 0.12 ± 0.68 | 1.00 | 35 | 34 |
| #00ff00 | #00ffa9 | 1.00 ± 0.49 | 0.91 ± 0.53 | 0.773 | 35 | 35 |
| #00ff00 | #00a9ff | 3.43 ± 1.20 | 3.34 ± 1.03 | 1.00 | 35 | 35 |
| #00ff00 | #0000ff | 3.83 ± 1.23 | 3.59 ± 1.12 | 0.751 | 35 | 35 |
| #00ff00 | #aa00ff | 5.57 ± 0.89 | 5.43 ± 1.03 | .884 | 35 | 35 |
| #00ff00 | #ff00aa | 5.77 ± 1.04 | 5.69 ± 0.99 | 1.00 | 35 | 35 |
| #00ffa9 | #00ffa9 | 0.18 ± 0.51 | 0.00 ± 0.00 | 0.511 | 34 | 33 |
| #00ffa9 | #00a9ff | 2.16 ± 0.64 | 2.39 ± 0.83 | 0.360 | 35 | 35 |
| #00ffa9 | #0000ff | 3.10 ± 1.09 | 3.31 ± 0.91 | 0.560 | 35 | 35 |
| #00ffa9 | #aa00ff | 4.89 ± 1.15 | 4.84 ± 1.08 | 1.00 | 35 | 34 |
| #00ffa9 | #ff00aa | 5.51 ± 1.11 | 5.90 ± 0.88 | 0.349 | 35 | 35 |
| #00a9ff | #00a9ff | 0.15 ± 0.43 | 0.24 ± 1.19 | 1.00 | 34 | 34 |
| #00a9ff | #0000ff | 1.2 ± 0.47 | 1.29 ± 0.62 | 1.00 | 35 | 35 |
| #00a9ff | #aa00ff | 2.89 ± 1.25 | 3.43 ± 1.2 | 0.071 | 35 | 35 |
| #00a9ff | #ff00aa | 5.91 ± 0.95 | 5.90 ± 0.87 | 1.00 | 35 | 35 |
| #0000ff | #0000ff | 0.00 ± 0.00 | 0.03 ± 0.17 | 1.00 | 35 | 34 |
| #0000ff | #aa00ff | 2.26 ± 0.96 | 2.47 ± 0.99 | 0.492 | 35 | 35 |
| #0000ff | #ff00aa | 6.09 ± 0.99 | 6.04 ± 0.98 | 1.00 | 35 | 35 |
| #aa00ff | #aa00ff | 0.03 ± 0.17 | 0.06 ± 0.33 | 1.00 | 34 | 35 |
| #aa00ff | #ff00aa | 1.89 ± 0.87 | 2.34 ± 0.99 | 0.112 | 35 | 35 |
| #ff00aa | #ff00aa | 0.26 ± 0.94 | 0.11 ± 0.67 | 0.970 | 35 | 35 |

##### 2-1-3. Intra-participant double-pass correlation analysis (Figure 3a diagonal)

To gauge the consistency of similarity ratings within individual participants, we conducted a double-pass correlation analysis on the similarity ratings pertaining to the 45 color pairs. This analysis focused on assessing the concordance between the first and second halves within participants, as illustrated in the diagonal of Figure 3a. The resulting histogram of correlation coefficients, depicted in Figure 3d, highlights a substantial level of coherence in ratings between the initial and subsequent halves.

**Figure 3.**
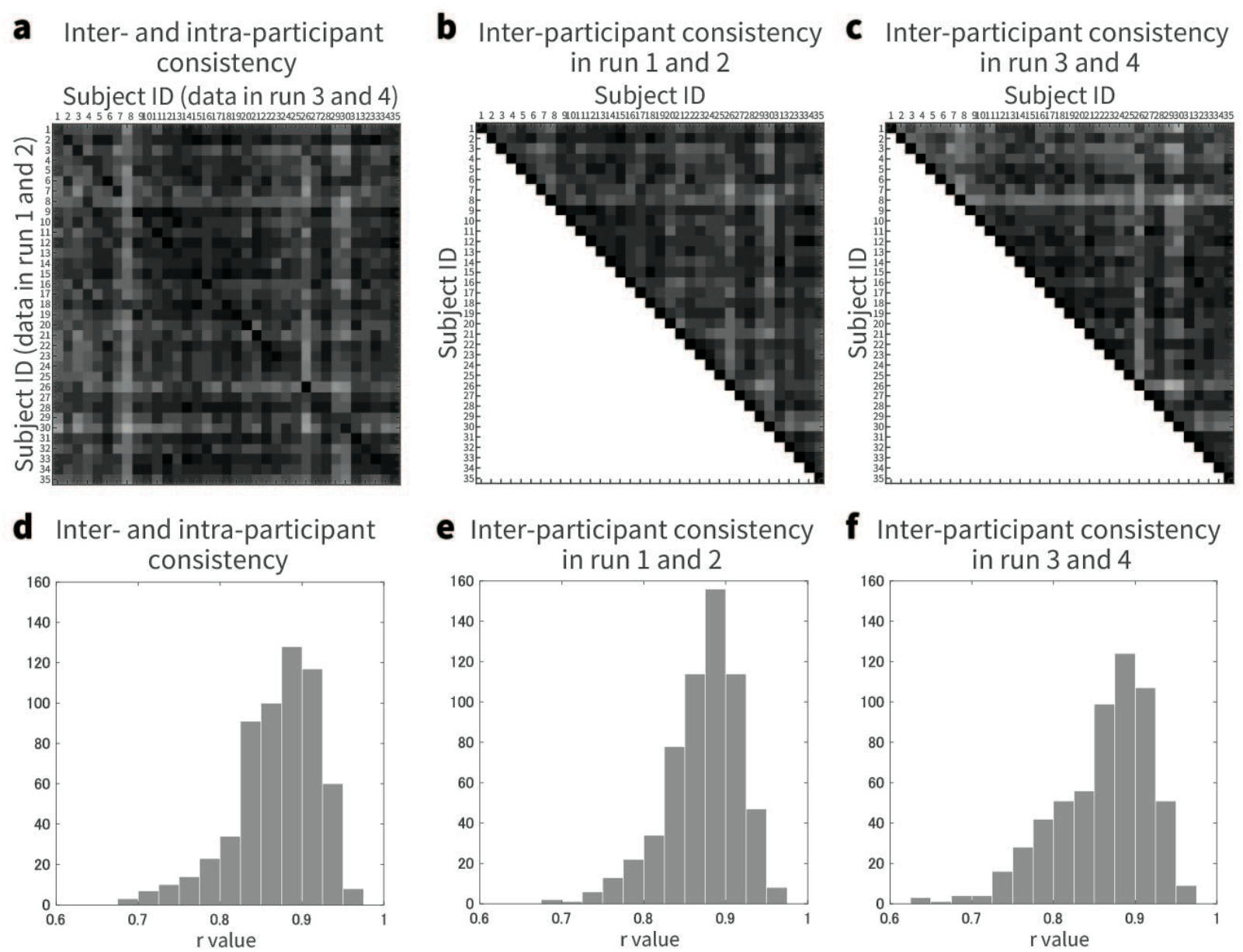
Analysis of inter- and intra-participant consistency in color pair evaluations. The figure presents correlation matrices that quantify intersubjective consistency across 45 color pairs among participants, with participant IDs labeled on both the X- and Y-axes. Each matrix element represents the correlation coefficient (r_i_j), which quantifies the consistency between the responses of participants i and j to the 45 color pairs. (a) The comparison of response patterns between runs 1-2 and runs 3-4, integrating intersubjective correlations within diagonal entries. (b) The intersubjective correlation of responses exclusively between run 1-2 and run 1-2. (c) The intersubjective correlation of responses exclusively between run 3-4 and run 3-4. (d-f) The histograms of correlation coefficients.

**Figure 4.**
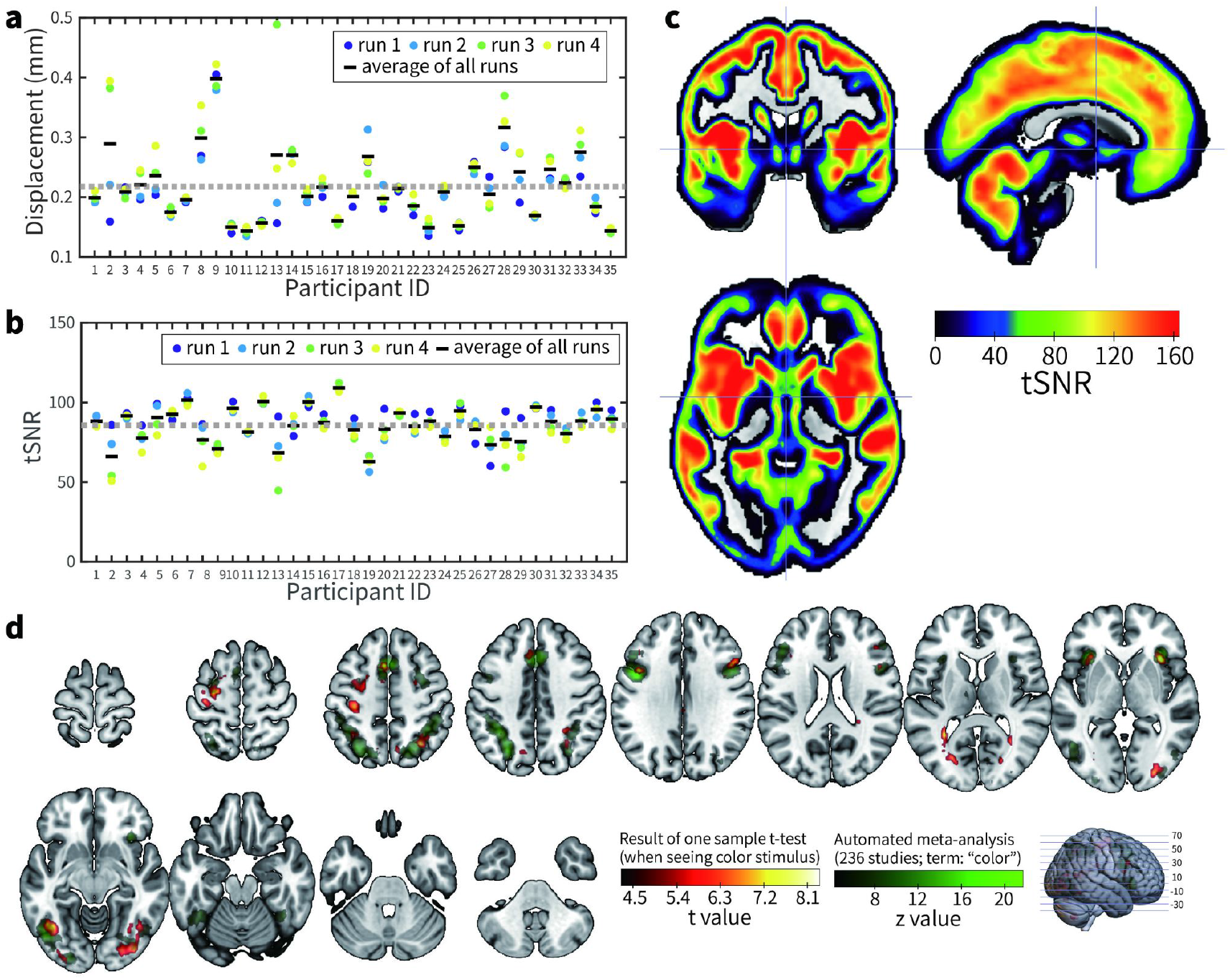
Visualization of fMRI-derived motion metrics, signal quality, and color observation-related brain activation. (a) Individual framewise displacement (FD data, detailing the average displacement per run and the overall average across all four runs for each participant. The gray dotted line shows the grand average with respect to the mean of all runs. (b) Individual data of tSNR. the average per run and the average of all four runs are shown for each individual. The gray dotted line represents the grand average for the mean of all runs. (c) Brain map for the voxel-wise tSNR. For visualization, tSNR was calculated using the data after the normalization process. (d) Activations for color observation. The result of the automated meta-analysis in the “color” term in Neurosynth is shown in green, and the activity during color observation in the present study is shown in hot color.

##### 2-1-4 Inter-participant correlation analysis (Figures 3a off diagonal, 3b, and 3c)

To evaluate the similarity of ratings across participants for the 45 color pairs, we computed correlations among the similarity ratings. Correlations were calculated for the first and second halves between participants (Figure 3a off-diagonal), as well as for the first half only (Figure 3b) and the second half only (Figure 3c).

Histograms illustrating the correlation coefficients resulting from these analyses are presented in Figures 3e and 3f. Notably, the correlation coefficients between participants were found to be consistently high, indicating a robust level of agreement in ratings across diverse participants.

#### 2.2 fMRI results

##### 2.2.1 FD and tSNR

FIgure 3b depicts the run-wise and across-run average of FD values. The across-run average FD of all 35 participants was 0.22 (SD: 0.06).

The average tSNR of all 35 participants was 85.77 (SD:10.68). These values were almost comparable to the typical values reported in previous fMRI data (A. Sengupta et al., 2016; The results of the voxel-wise tSNR are depicted in Figure 3d.

##### 2.2.2. Activation for color observation

Four participants with sudden and large body movements were excluded from the fMRI analysis. To confirm the brain activity during color observation, we calculated the activity when the target color stimulus was viewed without considering the type of color. Figure 3e shows the results of the one-sample t-test during the observation of the target color. The uniformity test map of the automated meta-analysis of 236 studies extracted with the word “color” in Neurosynth (https://neurosynth.org) was superimposed on the result of the one-sample t-test in the current study. A comparison of the Neurosynth results with the present results showed that the activation regions obtained in the present study were similar to the results of other color-related fMRI studies (e.g., visual areas and insular cortex).

## Code Availability

Other scripts used for technical validation in this study can be provided by the authors upon request. Because the color similarity rating task was programmed using Presentation software (Neurobehavioral Systems, Inc., Berkeley, CA), a software license is required to run the task. The other program codes, including those used in the analysis, were implemented in MATLAB.

## Acknowledgements

The authors would like to thank Sachie Sekimoto, Naoko Omorege, Wang Yaru, Mizuki Mori, and Ryo Michishita for their help with the data collection and organization. They also thank the staff of the Radiological Technology Section in the QST hospital for the acquisition of MRI data and Dr. Yoko Mizogami for her helpful comments on the construction of an experimental environment for color perception. This work was supported by JSPS KAKENHI

Grant (20H05710, 20H05711, 23H04830, 23H04833, 22H01108, 22K18265, 23K22379), JST Moonshot R&D Grant (JPMJMS2295-01, JPMJMS2295-02), JST CREST Grant (JPMJCR23P4) and MEXT Quantum Leap Flagship Program (MEXT QLEAP) Grant (JPMXS0120330644).

## Author contributions

T.H., D.M. and M.Y. drafted the manuscript. T.H. and D.M. contributed to data analysis and writing of the technical validation. T.H., M.M., D.M., N.T., and M.Y. designed the study. T.H., M.M., D.M., Y.T., T.O., and M.H. contributed to the data collection. T.H., D.M., and R.I. converted the dataset to the BIDS format. M.Y. supervised the project, and all authors reviewed the manuscript.

## Competing interests

The authors declare no competing interests.

## Notes

### Competing Interest Statement

The authors have declared no competing interest.

## References

1. D. C. Dennett, Quining Qualia, Consciousness in Modern Science, A. J. Marcel (eds.), E. Bisiach (ed.), Oxford University Press, (1988). D. Rosenthal, Quality Spaces and Sensory Modalities, Phenomenal Qualities: Sense, Perception, and Consciousness, Paul Coates (ed.), Sam Coleman (ed.), doi:10.1093/acprof:oso/9780198712718.003.0002, (2015).

2. S. B. Fink, L. Kob, H. Lyre, A structural constraint on neural correlates of consciousness, Philosophy and the Mind Sciences, 2, 7, doi:10.33735/phimisci.2021.79 (2021).

3. N. Tsuchiya and H. Saigo, A relational approach to consciousness: categories of level and contents of consciousness, Neuroscience of Consciousness, Volume 2021, 2, 2021, niab034, doi:10.1093/nc/niab034 (2021).

4. J. Kleiner, Mathematical Models of Consciousness, entropy, 22(6), 609, doi: 10.3390/e22060609

5. D. J. Chalmers, The Conscious Mind: In Search of a Fundamental Theory, Oxford University Press, (1996).

6. T. Nagel, What is it like to be a bat?, The philosophical review, 83, 4, 435–450, (1974).

7. N. Tsuchiya et al., Enriched category as a model of qualia structure based on similarity judgements, Consciousness and Cognition, Volume 101, doi:10.1016/j.concog.2022.103319, (2022).

8. C. E. Helm, Multidimensional Ratio Scaling Analysis of Perceived Color Relations, Journal of the Optical Society of America, 54, 256–262, doi:10.1364/JOSA.54.000256, (1964).

9. R. N. Shepard and L. A. Cooper, Representation of Colors in the Blind, Color-Blind, and Normally Sighted, Association for psychological science, Volume 3, 2, doi:10.1111/j.1467-9280.1992.tb00006.x, (1992)

10. G. P. Epping, E. L. Fisher, A. M. Zeleznikow-johnston, E. M. Pothos, N. Tsuchiya, A Quantum Geometric Framework for Modeling Color Similarity Judgments, Cognitive Science, Volume47, 1, doi:10.1111/cogs.13231, (2023).

11. Zeleznikow-Johnston, Y. Aizawa, M. Yamada, N. Tsuchiya, Are Color Experiences the Same across the Visual Field?, Journal of Cognitive Neuroscience, Volume 35, 4, doi:10.1162/jocn_a_01962, (2023).

12. T. Indow, Multidimensional Studies of Munsell Color Solid, Psychological Review, Volume 95, 4, 456–470, doi:10.1037/0033-295X.95.4.456, (1988).

13. G. V. Paramei, D. L. Bimler, C. R. Cavonius, Color-vision variations represented in an individual-difference vector chart, Color research and application, Volume 26, S1, doi:10.1002/1520-6378(2001)26:1+<::AID-COL49>3.0.CO;2-Z, (2001).

14. E. Boehm, D. I. A. MacLeod, J. M. Boesten, Compensation for red-green contrast loss in anomalous trichromats, journal of vision, Volume 14, 13, doi:10.1167/14.13.19, (2014).

15. G. Kawakita, A. Zeleznikow-Johnston, K. Takeda, N. Tsuchiya, M. Oizumi, Is my “red” your “red”?: Unsupervised alignment of qualia structures via optimal transport, PsyArXiv Preprints, doi:10.31234/osf.io/h3pqm, (2023).

16. J. M. Boesten, J. D. Robinson, G. Jordan, J. D. Mollon, Multidimensional scaling reveals a color dimension unique to ‘color-deficient’ observers, Current Biology, Volume 15, 23, doi:10.1016/j.cub.2005.11.031, (2005).

17. L. M. Vaina, Functional segregation of color and motion processing in the human visual cortex: clinical evidence, Cerebral Cortex, Volume 4, 5, 555–572, doi:10.109e3/cercor/4.5.555, (1994)

18. S. E. Bouvier, S. Engl, Behavioral Deficits and Cortical Damage Loci in Cerebral Achromatopsia, 16, 183–191, Cerebral Cortex, doi:10.1093/cercor/bhi096, (2006).

19. N. Hadjikhani, A. K. Liu, A. M. Dale, P. Cavanagh, R. B. H. Tootell, Retinotopy and color sensitivity in human visual cortical area V8, Nat Neurosci, 1, 235–241, doi:10.1038/681, (1998).

20. S. Zeki, J. D. Watson, C. J. Lueck, K. J. Friston, C. Kennard, R. S. Frackowiak, A direct demonstration of functional specialization in human visual cortex, Nat Neurosci, Volume 11, 3, doi:10.1523/JNEUROSCI.11-03-00641.1991, (1991).

21. A. Brewer, J. Liu, A.R. Wada, B. A. Wadell, Visual field maps and stimulus selectivity in human ventral occipital cortex, Volume 8, 8, 1102–1109, (2005).

22. W. Roe, L. Chelazzi, C. E. Connor, B. R. Conway, I. Fujita, J. L. Gallant, H. Lu, W. Vanduffel, Toward a Unified Theory of Visual Area V4, Neuron, Volume 74, 1, doi:0.1016/j.neuron.2012.03.011, (2012).

23. M. S. Beauchamp, J. V. Haxby, J. E. Jennings, E. A. DeYoe, An fMRI Version of the Farnsworth–Munsell 100-Hue Test Reveals Multiple Color-selective Areas in Human Ventral Occipitotemporal Cortex, Cerebral Cortex, Volume 9, 3, 257–263, doi:10.1093/cercor/9.3.257, (1999).

24. R. Shapley, M. J. Hawken, Color in the Cortex: single- and double-opponent cells, Vision Research, Volume 51, 7, 701–717, doi:10.1016/j.visres.2011.02.012, (2011).

25. A. Rosenthal, S. R. Singh, K. L. Hermann, D. Pantazis, B. R. Conway, Color Space Geometry Uncovered with Magnetoencephalography, Current Biology, Volume 31, 3, doi:10.1016/j.cub.2020.10.062, (2021).

26. G. J. Brouwer, D. J. Heeger, Decoding and Reconstructing Color from Responses in Human Visual Cortex, Journal of Neuroscience, Volume 29, 44, doi:10.1523/JNEUROSCI.3577-09.2009, (2009).

27. G. J. Brouwer, D. J. Heeger, Categorical Clustering of the Neural Representation of Color, Journal of Neuroscience, Volume 33, 39, doi:10.1523/JNEUROSCI.2472-13.2013, (2013).

28. N. Tsuchiya, M. Wilke, S. Frässle, V. A. F. Lamme, No-Report Paradigms: Extracting the True Neural Correlates of Consciousness, Trends in Cognitive Sciences, Volume 19, 12, 757–770, doi:10.1016/j.tics.2015.10.002, (2015).

29. A. Farooqui, T. Manly, When attended and conscious perception deactivates fronto-parietal regions, Cortex, Volume 107, 166–179, doi:10.1016/j.cortex.2017.09.004, (2018).

30. S. Frässle, J. Sommer, A. Jansen, M. Naber, W. Einhäuser, Binocular Rivalry: Frontal Activity Relates to Introspection and Action But Not to Perception, Journal of Neuroscience, Volume 34, 5, doi:10.1523/JNEUROSCI.4403-13.2014, (2014).

31. T. Dellert, M. Müller-Bardorff, I. Schlossmacher, M. Pitts, D. Hofmann, M. Bruchmann, T. Straube, Dissociating the Neural Correlates of Consciousness and Task Relevance in Face Perception Using Simultaneous EEG-fMRI, Journal of Neuroscience, doi:10.1523/JNEUROSCI.2799-20.2021, (2021).

32. M. A. Cohen, K. Ortego, A. Kyroudis, M. Pitts, Distinguishing the Neural Correlates of Perceptual Awareness and Postperceptual Processing, Journal of Neuroscience, doi:10.1523/JNEUROSCI.0120-20.2020, (2020).

33. Sergent, M. Corazzol, G. Labouret, F. Stockart, M. Wexler, J. King, F. Meyniel, D. Pressnitzer, Bifurcation in brain dynamics reveals a signature of conscious processing independent of report, Nat Commun, Volume 12, 1149, doi:10.1038/s41467-021-21393-z, (2021).

34. P. Shafto, M. A. Pitts, Neural Signatures of Conscious Face Perception in an Inattentional Blindness Paradigm, Journal of Neuroscience, Volume 35, 31, doi:10.1523/JNEUROSCI.0145-15.2015, (2015).

35. Stokes, M., Anderson, M., Chandrasekar, S. & Motta, R. A. (1996, November). standard default color space for the internet -sRGB. https://www.w3.org/Graphics/Color/sRGB.html

36. E. Rowe, M. Garrido, N. Tsuchiya, Feedforward connectivity patterns from visual areas to the front of the brain contain information about sensory stimuli regardless of awareness or report, PsyArXiv, doi:10.31234/osf.io/u36h8, (2023).

37. S. Ishihara, in Tokyo, Japan (ed 38 Plates Edition) (Kanehara & Co., 1990).

38. Japan Color Enterprise Co., LTD., Handbook for 100 hue test, Japan Color Enterprise Co., LTD.

39. R. C. Oldfield, The assessment and analysis of handedness: The Edinburgh inventory. Neuropsychologia, Volume 9, 1, doi:10.1016/0028-3932(71)90067-4, (1971).

40. D. Spielberger, F. Gonzalez-Reigosa, A. Martinez-Urrutia, L. F. Natalicio, D. S. Natalicio, The state-trait anxiety inventory. Revista Interamericana de Psicologia/Interamerican Journal of Psychology, Volume 5, 3, doi:10.30849/rip/ijp.v5i3%20&%204.620, (1971).

41. D. Spielberger, R. L. Gorsuch, R. E. Lushene., P. R. Vagg, G. A. Jacobs, Manual for the state-trait anxiety inventory, Palo Alto: Consulting Psychologists Press., (1983).

42. Beck, R. Steer, G. Brown, Manual for the Beck Depression Inventory-II. Psychological Corporation, (1996).

43. M. Kojima, T. A. Furukawa, H. Takahashi, M. Kawai, T. Nagaya, S. Tokudome, Cross-cultural validation of the Beck Depression Inventory-II in Japan. Psychiatry Research, Volume 110, 3, 291–299, doi:10.1016/s0165-1781(02)00106-3, (2002).

44. S. Moeller, E. Yacoub, C. A. Olman, E. Auerbach, J. Strupp, N. Harel, K. Uğurbil, Multiband multislice GE-EPI at 7 tesla, with 16-fold acceleration using partial parallel imaging with application to high spatial and temporal whole-brain fMRI. Magnetic Resonance in Medicine, Volume 63, 5, 1144–1153, doi:10.1002/mrm.22361, (2010).

45. D. Power, K. A. Barnes, A. Z. Snyder, B. L. Schlaggar, S. E. Petersen, Spurious but systematic correlations in functional connectivity MRI networks arise from subject motion. Neuroimage, Volume 59, 3, 2142–2154, doi:10.1016/j.neuroimage.2011.10.018, (2012).

46. J. D. Power, A. Mitra, T. O. Laumann, A. Z. Snyder, B. L. Schlaggar, S. E. Petersen, Methods to detect, characterize, and remove motion artifact in resting state fMRI. Neuroimage, Volume 84, 320–341, doi:10.1016/j.neuroimage.2013.08.048, (2014).

47. Sengupta, F. R. Kaule, J. S. Guntupalli, M. B. Hoffmann, C. Häusler, J. Stadler, M. Hanke, A studyforrest extension, retinotopic mapping and localization of higher visual areas. Sci Data, Volume 3, 160093, doi:10.1038/sdata.2016.93, (2016).

48. J. Berezutskaya, M. J. Vansteensel, E. J. Aarnoutse, Z. V. Freudenburg, G. Piantoni, M. P. Branco, N. F. Ramsey, Open multimodal iEEG-fMRI dataset from naturalistic stimulation with a short audiovisual film. Scientific Data, Volume 9, 91, doi:10.1038/s41597-022-01173-0. (2022).

